# Transcription repression of Cry2 via Per2 interaction promotes adipogenesis

**DOI:** 10.1101/2023.03.12.532323

**Authors:** Weini Li, Xuekai Xiong, Tali Kiperman, Ke Ma

## Abstract

The circadian clock is driven by a transcriptional-translational feedback loop, and Cryptochrome 2 (Cry2) represses CLOCK/Bmal1-induced transcription activation. Despite the established role of clock in adipogenic regulation, whether the Cry2 repressor activity functions in adipocyte biology remains unclear. Here we identify a critical cysteine residue of Cry2 that mediates interaction with Per2, and demonstrate that this mechanism is required for clock transcriptional repression that inhibits Wnt signaling to promote adipogenesis. Cry2 protein is enriched in white adipose depots and was robustly induced by adipocyte differentiation. Via site-directed mutagenesis, we identified that a conserved Cry2 Cysteine at 432 within the loop interfacing with Per2 mediates heterodimer complex formation that confers transcription repression. C432 mutation disrupted Per2 association without affecting Bmal1 binding, leading to loss of repression of clock transcription activation. In preadipocytes, whereas Cry2 enhanced adipogenic differentiation, the repression-defective C432 mutant suppressed this process. Furthermore, silencing of Cry2 attenuated, while stabilization of Cry2 by KL001 markedly augmented adipocyte maturation. Mechanistically, we show that transcriptional repression of Wnt pathway components underlies Cry2 modulation of adipogenesis. Collectively, our findings elucidate a Cry2-mediated repression mechanism that promotes adipocyte development, and implicate its potential as a clock intervention target for obesity.

## Introduction

The circadian clock, a molecular machinery that generates daily oscillations in behavior and physiology, is driven by a transcriptional/translational feed-back loop (Takahashi 2017). This pervasive temporal mechanism orchestrates rhythmic regulations of metabolic processes that is required to maintain homeostasis (Panda et al. 2002; Xiong et al. 2021). Key transcription activators of the core clock loop, CLOCK (Circadian Locomotor Output Cycles Kaput) and BMAL1, initiate transcription of their target genes to drive clock oscillation (Gekakis et al. 1998; Huang et al. 2012). CLOCK/BMAL1-controlled transcription activation leads to expression of clock repressors, the Periods (Per1-3) and Cryptochromes (Cry1 & 2) genes, that exert negative control of CLOCK/BMAL1 activity in gene transcription (Shearman et al. 2000). The PER and CRY proteins block the transcriptional activity of CLOCK:BMAL1 by forming a repressor complex that constitutes the negative feedback arm of the core clock regulatory mechanism (Nangle et al. 2014; Crosby and Partch 2020).

The CRY repressor proteins are essential components of the circadian timekeeping machinery required for clock function (van der Horst et al. 1999; Vitaterna et al. 1999). Posttranslational mechanisms involved in modulating CRY stability and activity are integral determinants of circadian oscillation. Particularly, the spatiotemporal regulation of its interaction with Per proteins drives the PER-CRY repressive complex and transcription repression of CLOCK:BMAL1 (Xing et al. 2013; Hirano et al. 2014; Nangle et al. 2014; Schmalen et al. 2014). The abundant and post-translational modifications of Per is critical for the formation of PERs-CRY complex and nuclear localization (Takano et al. 2004; Reischl et al. 2007; Chen et al. 2009; Ye et al. 2011; St John et al. 2014; Aryal et al. 2017). The F-box ubiquitin E3 ligases, β-TrCP and FBXL3, modulate PER and CRY degradation, respectively, to control circadian protein stability (Reischl et al. 2007; Siepka et al. 2007; Xing et al. 2013). Interaction of PER with CRY to form a repression complex thus prevent CRY from E3 ligase FBXL3-medaited degradation (Busino et al. 2007; Godinho et al. 2007; Siepka et al. 2007; Nangle et al. 2014). Interestingly, an interface loop within the Cry2 FAD binding domain was identified to form a zinc finger structure required for interaction with Per1/2 (Nangle et al. 2014; Schmalen et al. 2014), and conserved residues of this interface are also involved in Cry2 binding activity with FBXL3. A recently identified Cry-stabilizing molecule, KL001, functions via a similar action by blocking FBLX3 interaction with CRY2 to prevent ubiquitination- mediated degradation that promotes CRY2 protein stability (Hirota et al. 2012; Lee et al. 2015).

The circadian clock exerts temporal control in adipocyte development (Bass and Takahashi 2010; Takahashi 2017), and key components of the molecular clock are involved in modulating adipocyte maturation (Nam et al. 2016). A daily burst of commitment to differentiation occurs in preadipocytes driven by accumulation of adipogenic factors occurs within a 12-hour window (Zhang ZB et al. 2022). The essential clock activator Bmal1 inhibits adipogenesis via direct transcriptional control of signaling components involved in Wnt pathway, a potent developmental signaling that suppresses adipocyte development with a similar effect on beige adipocyte development (Guo et al. 2012; Xiong et al. 2022). Rev-erbα, a direct clock repressor target of Bmal1 that functions as the negative arm of the re-enforcing loop of the clock regulatory network, modulates brown adipogenesis. Cry proteins are involved in metabolic regulations. Cry1 antagonizes cAMP-mediated hepatic gluconeogenesis in liver (Zhang EE et al. 2010), while impaired Cry1 degradation contributes to hyperglycemia in diabetic models (Kim et al. 2022). On the other hand, compounds that stabilize Cry2 protein against proteasome-mediated degradation, KL001 and its derivatives, demonstrate anti-gluconeogenic effects with glucose-lowering activity of the derivative *in vivo* (Humphries et al. 2016, 2018). Thus, modulation of Cry2 activity may have potential applications for metabolic disease therapy.

Despite the role of molecular clock modulation in adipocyte development, whether the clock negative feedback mechanism *via* Cry-Per interaction is involved in adipogenic regulations remains unknown. In the current study, we employed distinct approaches to probe the transcription repressor function of Cry2 and its interaction with Per2 to define how a Cry2-mediated clock inhibitory mechanism modulates adipocyte maturation.

### Materials & Methods

#### Animals

Mice were maintained in the City of Hope vivarium under a constant 12:12 light dark cycle, with lights on at 6:00 AM (Zeitgeber Time ZT0). All animal experiments were approved by the Institutional Animal Care & Use Committee (IACUC) of City of Hope and preformed according to approval. C57BL/6J mice were purchased from Jackson Laboratory and used for experiments following 2 weeks or longer of acclimation.

#### Cell culture and reagents

Pre-adipocyte 3T3-L1 (CL-173) and L Wnt3A-expressing cells (CRL-2647) were purchased from the American Type Culture Collection (ATCC). 3T3-L1, Wnt3A and HEK293A cells were cultured in Dulbecco’s Modified Eagle Medium (Gibco) supplemented with 10% Fetal bovine serum (FBS, Cytiva) and Penicillin-Streptomycin-Glutamine (PSG, Gibco). Wnt3a-conditioned media was collected from Wnt3a-producing L cell using the ATCC standard protocol and diluted in a 1:10 ratio or at the concentration indicated. The transient transfection reagents used were from Lipofectamine 3000 (Invitrogen), Polyethylenimine MAX (PEI MAX) (Polysciences) or PolyJet (SignaGen Laboratory). The chemical reagents used include Puromycin (Sigma Aldrich), Cycloheximide and Polybrene (Santa Cruz), and KL001 (Cayman Chemicals).

#### Plasmids and plasmid construction

Bmal1-His was a gift from Aziz Sancar (Addgene plasmid #31367)(Ye et al. 2011). Flag-HA-GFP was a gift from Wade Harper (Addgene plasmid # 22612)(Sowa et al. 2009). PGL2 basic vector were obtained from Addgene. PGL2 basic vector was a gift from Joseph Takahashi (Addgene plasmid # 48742)(Yoo et al. 2005). Full-length CLOCK and full-length Per2 were generated from mouse cDNA and cloned into the PcDNA3.0-Myc vector. Full-length Cry2 was constructed by subcloning the corresponding cDNA into PcDNA3.0-Flag or pCDH-puro vector. Cry2 cysteine mutations (Cry2 C430A, Cry2 C432A and Cry2 C430A/C432A) were constructed via site-specific mutagenesis and cloned into PcDNA3.0-Flag or pCDH- puro vector. The primer sequences used for mutagenesis were listed in Supplementary Table S1. The lentiviral shRNA sequences for mouse Cry2 were subcloned into the pLKO-puro vector and the sequences were listed in Supplementary Table S2.

#### Lentivirus packaging and infection for generation of stable cell line

The 293A human embryonic kidney cells were transfected with the packaging plasmids (pSPAX.2 and pMD2.G) and recombinant lentivirus vectors using the PEI reagent according to the manufacturer’s protocol. After 48h post-transfection, lentiviruses were collected through 0.45 μm filter to remove the cell debris. The 3T3-L1 cell lines were infected by the lentivirus medium supplemented with 8 μg/ml polybrene. After the 24h infection, stable cell lines were selected in the presence of 1 μg/ml or 2 μg/ml puromycin for selection of stable clones. The stable cell lines overexpressing or knocking down Cry2 were verified by RT-qPCR or immunoblotting.

#### Luciferase Assay

The luciferase reporter assay was performed according to manufacturer’s protocol for the Dual- Luciferase® Reporter Assay Kit (Promega), as previously described(Guo et al. 2012). Briefly, cells were seeded on 24 well plates and transfected with luciferase reporter and indicated plasmids. After 24h transfection, cells were extracted with lysis buffer for 30 min and transferred to 96-well plate. LARII and Stop & Glo components were added for measuring luciferase activity. Bioluminescence was measured using a microplate reader (TECAN infinite M200pro). The mean and standard deviation values relative to Renilla were calculated for each well for 4 wells and graphed.

#### Immunoprecipitation and immunoblot analysis

The HEK 293A were transfected with the indicator plasmids and lysed in the IP buffer containing 50 mM Tris–HCl pH 7.4, 150 mM NaCl, 1% TritonX-100 and 10% glycerol supplemented with protease inhibitor (Thermo Fisher Scientific) for 30mins. The supernatants were collected and incubated with anti-Flag magnetic beads (Sigma Aldrich), anti-Myc agarose beads (Thermo Fisher) or antibodies containing protein A/G beads at 4 °C overnight. The complex was washed with IP buffer three times followed by immunoblotting analysis. Total protein (20-30 μg) samples were loaded on 10% SDS-PAGE gels and transferred to PVDF membranes (Bio-rad). NC membrane was blocked with 5% nonfat dry milk or FBS in PBS with 0.1% Tween 20 (PBST) for one hour at RT and incubated with primary antibodies at 4°C overnight. Primary antibodies used were listed in Suppl. Table S3. After washing three times in PBST, the membrane incubated with secondary antibodies for one hour. The images were developed using the chemiluminescence imager (GE Healthcare BioSciences AB Amersham Imager 680) and analyzed by Image J.

#### RNA extraction Real-time PCR

Total RNA was isolated from culture cells or tissues using Trizol reagent (Thermo Fisher). 2 μg RNA was converted to cDNA using Revert-Aid RT Reverse Transcription Kit (Thermo Fisher). The Real-time RCR was performed on ViiA 7 Real-Time PCR system (Applied Biosystems) with PowerUp SYBR Green Master Mix (Thermo Fisher) according to the manufacturer’s protocol. The sequences of primers were listed in Suppl. Table S4.

#### Primary preadipocytes isolation and adipogenic induction

The stromal vascular fraction containing preadipocytes were isolated from subcutaneous fat pads, as described (Chatterjee et al. 2013). Briefly, fat pads were cut into pieces and digested using 0.1% collagenase Type 1 with 0.8% BSA at 37^0^C in a horizontal shaker for 60 minutes, passed through Nylon mesh and centrifuged to collect the pellet containing the stromal vascular fraction with preadipocytes. Preadipocytes were cultured in F12/DMEM supplemented with bFGF (2.5 ng/ml), expanded for two passages and subjected to differentiation in 6-well plates at 90% confluency. Adipogenic differentiation was induced for 2 days in medium containing 10% FBS, 1.6 μM insulin, 1 μM dexamethasone, 0.5 mM IBMX before switching to maintenance medium for 4 days with insulin only.

#### Oil-Red-O and Bodipy 493/503 staining of lipids droplets

Staining of oil-Red-O and Bodipy 493/503 were performed as previously described (Guo et al. 2012; Xiong et al. 2022). Briefly, 3T3-L1 were induced to differentiation at indicated duration, washed gently three times with PBS and fixed using 10% formalin. For oil-Red-O staining, the fixed cells were washed twice with ddH2O and then dried for 30 minutes at room temperature. Cells were then incubated in 0.5% Oil-Red-O solution for 20 minutes and washed with ddH2O. For Bodipy staining, cells were fixed by 10% formalin for 30 min and permeabilized with 0.2 % TritonX-100 for 10 minutes. The cells were then washed using PBS and incubated using Bodipy 493/503(1:1000) and DAPI (1:2000) for 20 minutes.

#### TOPFlash luciferase reporter assay

M50 Super 8x TOPFlash luciferase reporter containing Wnt-responsive TCF bindings sites was a gift from Randall Moon (Addgene #12456)(Veeman et al. 2003). Cells were seeded at 4x10^5^ concentration in 24-well plates and transfection was performed at 90% confluence. 24 hours following transfection, cells were treated using 10% Wnt3a media to induce luciferase activity. Luciferase activity was assayed using the Dual-Luciferase Reporter Assay Kit (Promega) in 96-well black plates 24 hours after Wnt3a stimulation, as previously described(Chatterjee et al. 2013). The luminescence was normalized to Renilla and the mean and standard deviation values from 4 wells were graphed.

#### Statistical analysis

Data from minimum of three independent experiments were shown as mean ± SD or SEM as indicated. Sample size were indicated for each experiment in figure legends. Each experiment was repeated at minimum twice to validate the result. Two-tailed Student’s t-test or one-way ANOVA with post-hoc analysis for multiple comparisons were performed as appropriate as indicated. P<0.05 was considered statistically significant.

### Results

#### Cry2 rhythmic expression is enriched in visceral adipose tissue depot and induced during adipocyte differentiation

Adipose tissue possesses cell-autonomous circadian clock (Zvonic et al. 2006; Wu et al. 2007). To study the potential function of Cry2 in adipocytes, we examined Cry2 protein rhythm in representative adipose depots. Interestingly, Cry2 was highly enriched in classic white adipose tissue (epidydimal eWAT) as compared to liver or muscle (Fig, 1A), or thermogenic fat depots including the inguinal fat (iWAT) or brown adipose tissue (BAT, Fig. 1B). Diurnal expression of Cry2 in adipose tissues examined was evident, with elevated levels observed at late night hour ZT23 (ZT: Zeitgieber Time). Further analysis of Cry2 protein rhythm in eWAT every four hours over one circadian cycle revealed its peak expression at near ZT18 (Fig. 1C). As expected, mRNA transcript of Cry2 and closely related Cry1 displayed circadian profiles peaking at ZT 20-24 in both eWAT and iWAT depots (Fig. 1D & 1E). To explore the potential role of Cry2 in adipogenesis, we examined its expression dynamic during adipogenic differentiation of 3T3-L1 preadipocytes. Cry2 transcript was robustly induced from day 0 to day 8 of differentiation, similar to that of the adipogenic factors *PPARγ* and *C/EBPα* (Fig. 1F). Cry2 protein level was also markedly elevated in differentiated adipocytes as compared to undifferentiated preadipocytes (Fig. 1G). Examination of Cry2 protein expression in primary preadipocytes isolated from subcutaneous fat further confirmed its robust induction in mature adipocytes than that of the adipogenic progenitors (Fig. 1H).

**Figure 1.**
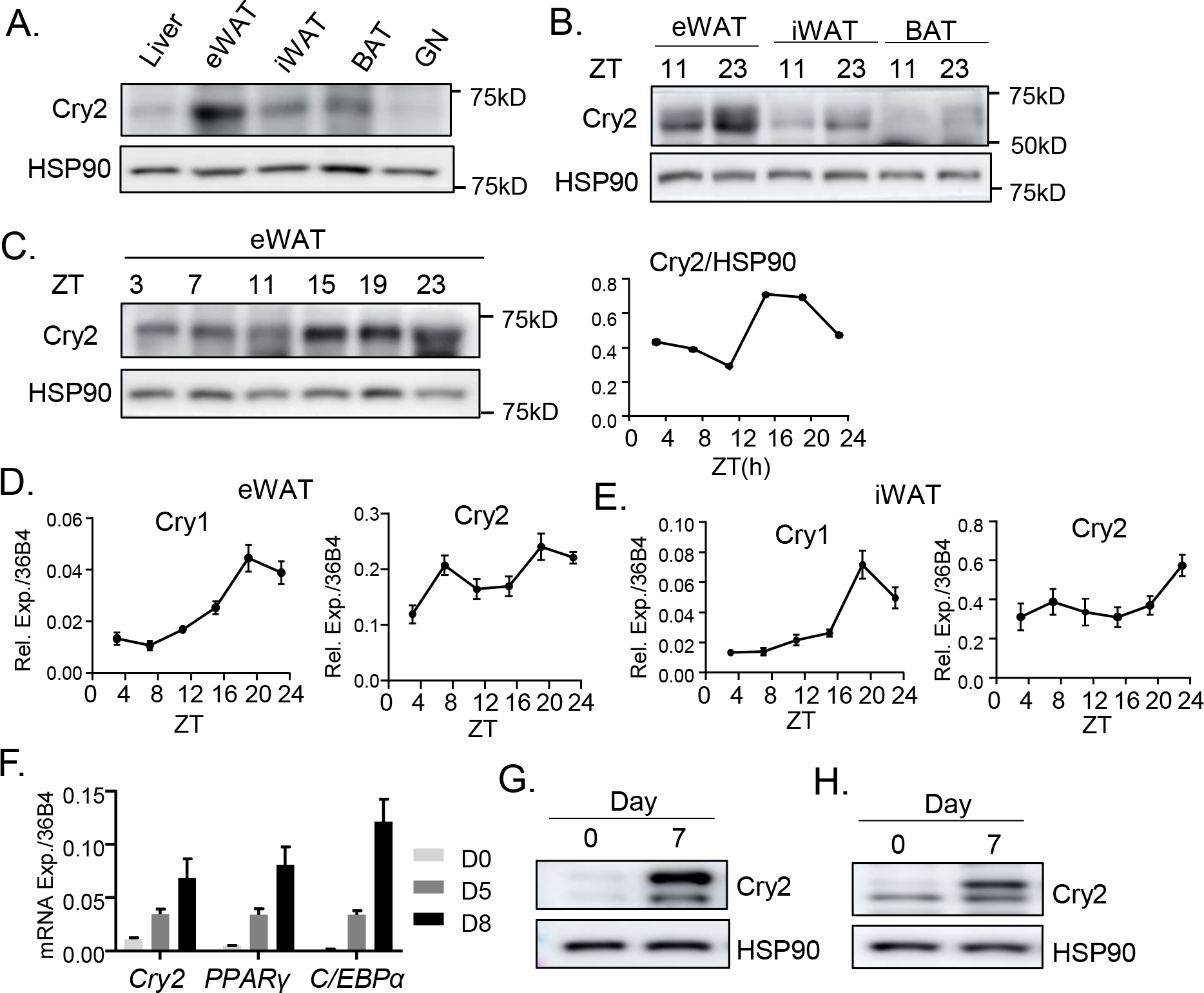
Cry2 is enriched in adipose depots. (A, B). Immunoblot analysis of Cry2 expression in liver, epidydimal white adipose tissue (eWAT), inguinal white adipose tissue (iWAT), brown adipose tissue (BAT) and gastrocnemius muscle (GN) at ZT3 (zeitgeber time with light on as ZT0, A), and its expression at ZT11 and ZT23 in distinct adipose depots (B). (C) Circadian protein profile of Cry2 at indicated zeitgeber times (ZT) every four hours over a 24-hour cycle. Each lane represents pooled samples of 4-5/time point. (D, E) RT-qPCR analysis of circadian transcript expression of Cry1 and Cry2 in eWAT (D) and iWAT (E). Expression were normalized to 36B4, and error bars represent SE of 5 mice per time point. (F) RT-qPCR analysis of Cry2 and adipogenic factors at 0, 5 and 8 days during 3T3-L1 differentiation. n=3/group. (G, H) Immunoblot analysis of Cry2 protein induction at day 0 and day 7 during adipogenic differentiation of 3T3-L1 (G), and primary preadipocytes (H).

#### A Cry2 Cysteine 432 mutation disrupts interaction with Per2 with loss of repressive activity

Cry2 functions in a transcription repressor complex with Per2 protein to inhibit CLOCK/Bmal1-mediated transcription activation (Gustafson and Partch 2015; Crosby and Partch 2020). The coiled-coil region of Cry2 was found to interact with Per2 and FBXL3, with Per2 binding blocking Cry2 degradation by preventing FBXL3-mediated ubiquitination (Nangle et al. 2014). Cysteine residues 430 and 432 are conserved homologous residues between Cry2 and Cry1 that lie within an interface loop between α-helixes 18 and 19 that may interact with Per2 (Fig. 2A). As dimerization with Per2 is required for Cry2 transcriptional repression, we tested whether these residues mediate Cry2-Per2 repression complex and thus the repressor activity of Cry2. Co-immunoprecipitation using pull-down of Flag-tagged Cry2 revealed significantly reduced binding with Myc-tagged Per2 when C432 was mutated to alanine, with even lower binding detected when both C432 and C430 were mutated, although the C430A mutant alone did not affect the interaction (Fig. 2B). The same result was obtained with these mutants using Myc-tag pull-down of Per2 protein, with nearly abolished Cry2-Per2 interaction of the C432A single and C430A/C432A double mutants but not C430A mutant (Fig. 2C), indicating that C432 is required for Cry2- Per2 heterodimer formation. We next tested whether this residue of Cry2 is involved in interaction with the CLOCK/Bmal1 activator complex. Co-immunoprecipitation with either Cry2 pull-down (Fig. 2D) or Bmal1 antibody (Fig. 2E) indicated that neither C432 nor C430 mutation attenuated Cry2 association with Bmal1, while C430A may modestly promote interaction between Cry2 with Bmal1. As Cry2-Per2 forms a repressor complex to inhibit CLOCK/Bmal1-mediated core clock transcription, we tested whether C432 mutation affects transcription repressor activity of Cry2 using a *Per2::dLuc* luciferase reporter containing canonical E-box CLOCK/Bmal1 binding sites(Yoo et al. 2004; Yoo et al. 2005). As expected, wild-type Cry2 completely suppressed CLOCK/Bmal1-activated transcription to basal level (Fig. 2F). In contrast, C432A mutant nearly abolished Cry2 repression of CLOCK/Bmal1 transcription activation that was largely comparable to vector control. Interestingly, while C430A/C432A double mutant displayed loss of repression activity similar to that of C432A, the C430A mutant enhanced Cry2-mediated repression. We next tested whether loss of repressor activity of the Cry2 C432A mutant is due to altered interaction with Fbxl3 affecting protein stability(Siepka et al. 2007) and found that this mutation instead abolished Cry2 binding with Fbxl3 (Fig. 2G). In addition, the loss of C432A mutant interaction with Fbxl3 was not altered by CRY2 stabilizing compound KL001 that functions by reducing Cry2 binding with Fbxl3(Hirota et al. 2012). Consistent with impaired Fbxl3 association, C432A mutant displayed improved stability as compared to wild-type Cry2 protein, as indicated by analysis of protein half-life following cycloheximide treatment (Fig. 2H).

**Figure 2.**
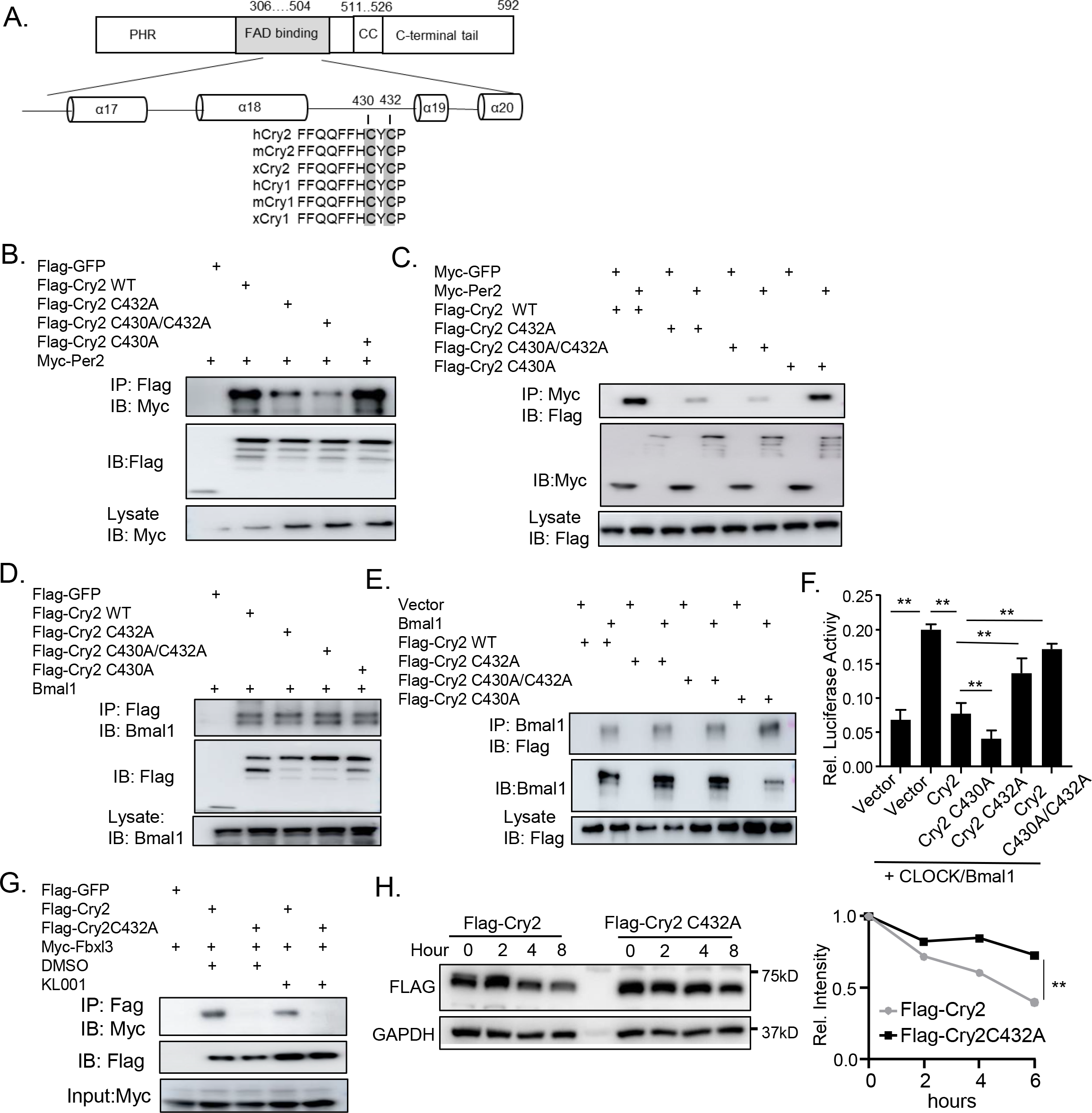
Mutation of Cry2 interface loop impairs interaction with Per2 and transcription repression. (A) Schematic presentation of Cry2 protein domain structure, with the conserved C430 and C432 within the interface loop region indicated. (B, C) Co-immunoprecipitation analysis of interaction of Flag-tagged Cry2 or C430/432 mutants with Myc-tagged Per2 protein. HEK293A cells were transfected with plasmids encoding Cry2, Cry2 mutants or Per2 and subjected to immunoprecipitation with anti-Flag antibody- conjugated agarose beads (B) or anti-Myc beads (C) with immunoblot analysis for Myc-tag (B) or Flag tag (C). (D, E) Co-immunoprecipitation analysis of interaction of Flag-tagged Cry2 or C430/432 mutants with Bmal1 protein. HEK293A cells were co-transfected with Bmal1 and Flag-tagged Cry2 or Cry2 mutants, subjected to immunoprecipitation with anti-flag beads (D), or anti-Bmal1 antibody (E) and followed by immunoblotting. (F) Analysis of transcription repression of Cry2 and the mutants on CLOCK/Bmal1-activated *Per2:dLuc* luciferase reporter. HEK293A were transiently transfected with *Per2:dLuc* luciferase reporter (40 ng), Renilla for normalization of transfection efficiency (10 ng), CLOCK, Bmal1 and indicated Cry2 expression plasmids (400 ng each). Error bars represent SD of four experimental repeats. (G) Co-immunoprecipitation analysis of interaction of Flag-tagged Cry2 or C432A mutant with Myc-tagged Fbxl3 protein. HEK293A cells were co-transfected with Flag-tagged Cry2 or Cry2 mutant and Myc-tagged Fbxl3, and subjected to immunoprecipitation with anti-Flag beads followed by immunoblotting using anti-Myc antibody. (H) Analysis of Cry2 and C432A mutant protein stability via determination of half-life following cycloheximide treatment (CHX). HEK293A were transfected with Flag-tagged Cry2 or C432A plasmids and were treated CHX 50 ng/mL at indicated time 24 hour after transfection, followed by immunoblot analysis. Densitometry quantification of protein abundance was normalized to GAPDH. **: P≤0.01 by one-way ANOVA.

#### Cry2 ectopic expression promoted and loss-of-function mutation abolished adipogenic differentiation

We previously demonstrated that transcription activator of the clock feedback loop Bmal1 inhibits adipogenesis (Guo et al. 2012). Given Cry2 repression of CLOCK/Bmal1-medaited transcription and the loss of this activity in the C432A mutant, Cry2 and this repression-defective mutant may exert opposing effects on adipogenic regulation. We thus generated 3T3-L1 preadipocytes with stable expression of Cry2, C432A and C430A/C432A mutants. Cry2 and the C432A mutants displayed comparable levels of protein expression (Fig. 3A). By subjecting to adipogenic differentiation via a standard induction cocktail(Guo et al. 2012; Xiong et al. 2022), cells with ectopic expression of wild-type Cry2 displayed markedly increased lipid accumulation indicative of enhanced differentiation efficiency compared to cells containing vector controls, as indicated by oil-red-O staining (Fig. 3B). In contrast, enhanced mature adipocyte formation was abolished in Cry2 C432A-expressing preadipocytes, with comparable extent of lipid staining as observed in controls. Fluorescence Bodipy staining for lipids further validated this finding (Fig. 3C). Analysis of adipocyte differentiation in Cry2-expressing cells revealed marked inductions of adipogenic factors C/EBPα and PPARγ, together with up-regulation of mature adipocyte marker FABP4 after 6 days near complete differentiation, while this effect was abolished in the C432A mutant (Fig. 3D). RT-qPCR analysis revealed similar findings of C/EBPα and FABP4 inductions by ectopic expression of Cry2 but not the C432A mutant, whereas C/EBPβ was not significantly altered (Fig. 3E). Furthermore, consistent with the stronger loss of repression activity of the C430A/C432A mutant, it resulted in a severe blockade of adipogenic differentiation that was lower than the controls as indicate by lipid staining via oil-red-O or Bodipy (Fig. 3F & 3G).

**Figure 3.**
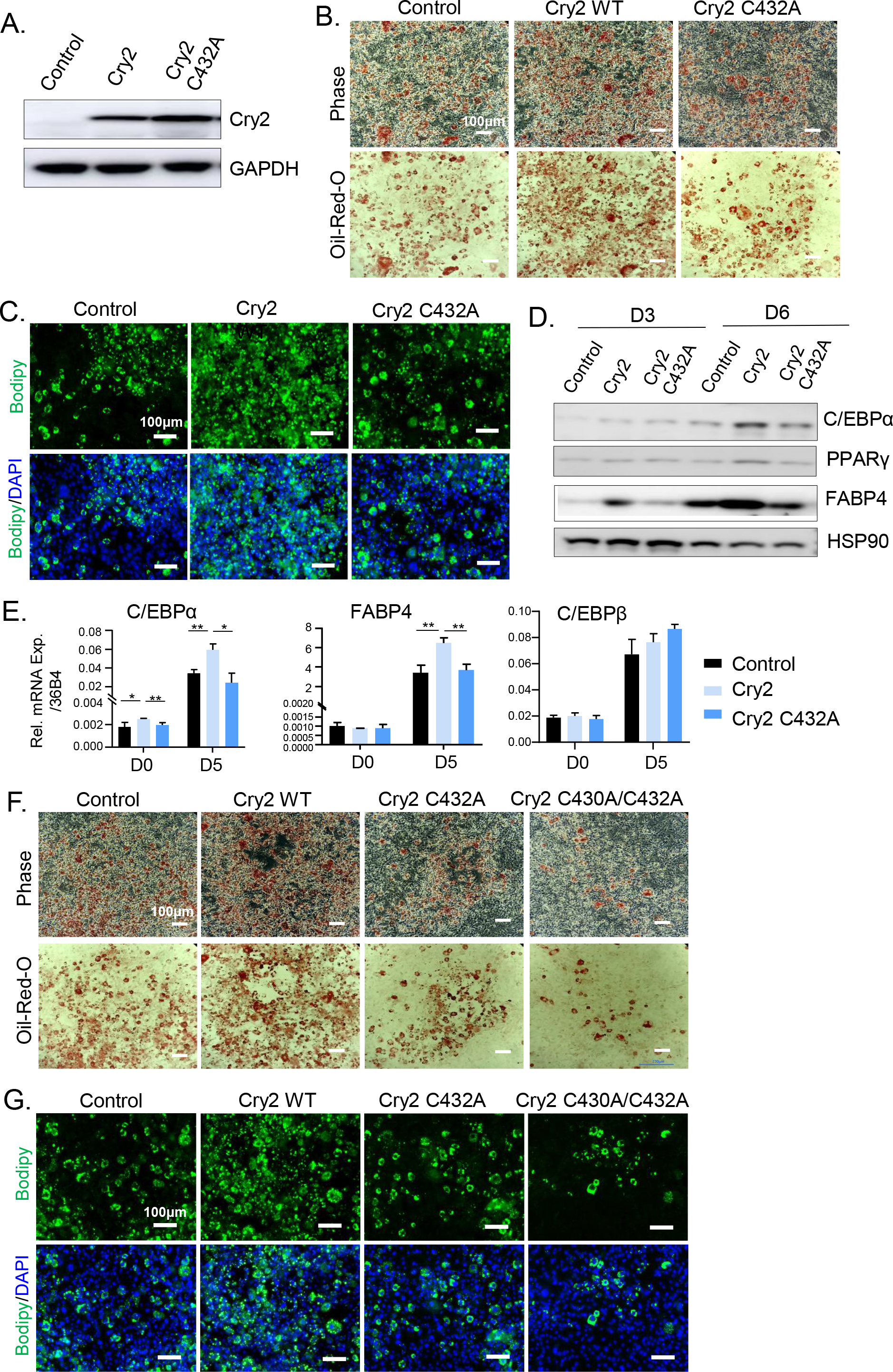
Cry2 promotes adipogenesis with loss of effect in repression-defective mutants. (A) Immunoblot analysis of Cry2 protein expression in 3T3-L1 stable clones overexpressing wild-type Cry2 or Cry2 C432A mutant. (B, C) Representative images of oil-red-O (B) and Bodipy staining (C) of 3T3-L1 with stable expression of Cry2 or C432A mutant at day 8 of adipogenic differentiation. (D, E) Immunoblot analysis (D), and RT-PCR analysis (E) of adipogenic and mature adipocyte markers at day 3 and day 6 (D) or day 5 of differentiation (E) of 3T3-L1 with ectopic expression of Cry2 or C432A mutant. (F, G) Representative images of oil-red-O (F) and Bodipy staining (G) of 3T3-L1 with stable expression of Cry2, C432A and C430A/C432A mutants at day 8 of adipogenic differentiation.

#### Cry2 exert transcriptional inhibition of Wnt signaling pathway

Clock exerts transcriptional control of key components of the Wnt signaling pathway leading to inhibition of adipogenesis (Guo et al. 2012). Cry2, as a clock repressor, may promote adipocyte differentiation via transcriptional repression of clock-controlled Wnt signaling. We therefore determined mRNA expression levels of Wnt pathway components that were direct transcriptional targets of CLOCK/Bmal1 (Guo et al. 2012). Forced expression of Cry2 significantly attenuated expressions of several Wnt signaling steps, including Wnt receptors Frizzed 2 (Fzd2), Dishevelled 2 (Dvl2), and the key signaling mediator β-catenin (Fig. 4A). In comparison, Cry2 C432A mutant induced Wnt1 and β-catenin expression as compared to control, suggesting opposite regulation of Wnt signaling activity with Cry2. Examination of total β-catenin protein level revealed marked reduction in Cry2-expressing preadipocytes most evident at day 5 of differentiation, indicating reduced Wnt activity (Fig. 4B). In contrast, Cry2 C432A mutant expression elevated total β-catenin protein as compared to Cry2-expressing cells, at a similar level as cells expressing vector control suggesting rescued Wnt signaling that reversed the suppressive effect by wild-type Cry2. To determine whether Cry2 inhibition of Wnt signaling mediates its effect on inducing adipogenesis, we treated Cry2-expressing preadipocytes with Wnt3a and found that it was sufficient to block Cry2-induced mature adipocyte formation. In control 3T3-L1 cells, Wnt3a completely abolished the mature adipocyte differentiation as indicated by the loss of lipid staining by Bodipy (Fig. 4C) or oil-red-O (Fig. 4D), as expected. Notably, Wnt3a was equally effective in preventing the enhanced adipogenic differentiation efficiency of Cry2-expressing cells, suggesting that the restoration of Wnt signaling by Wnt3a stimulation abolished Cry2 effect on adipogenesis.

**Figure 4.**
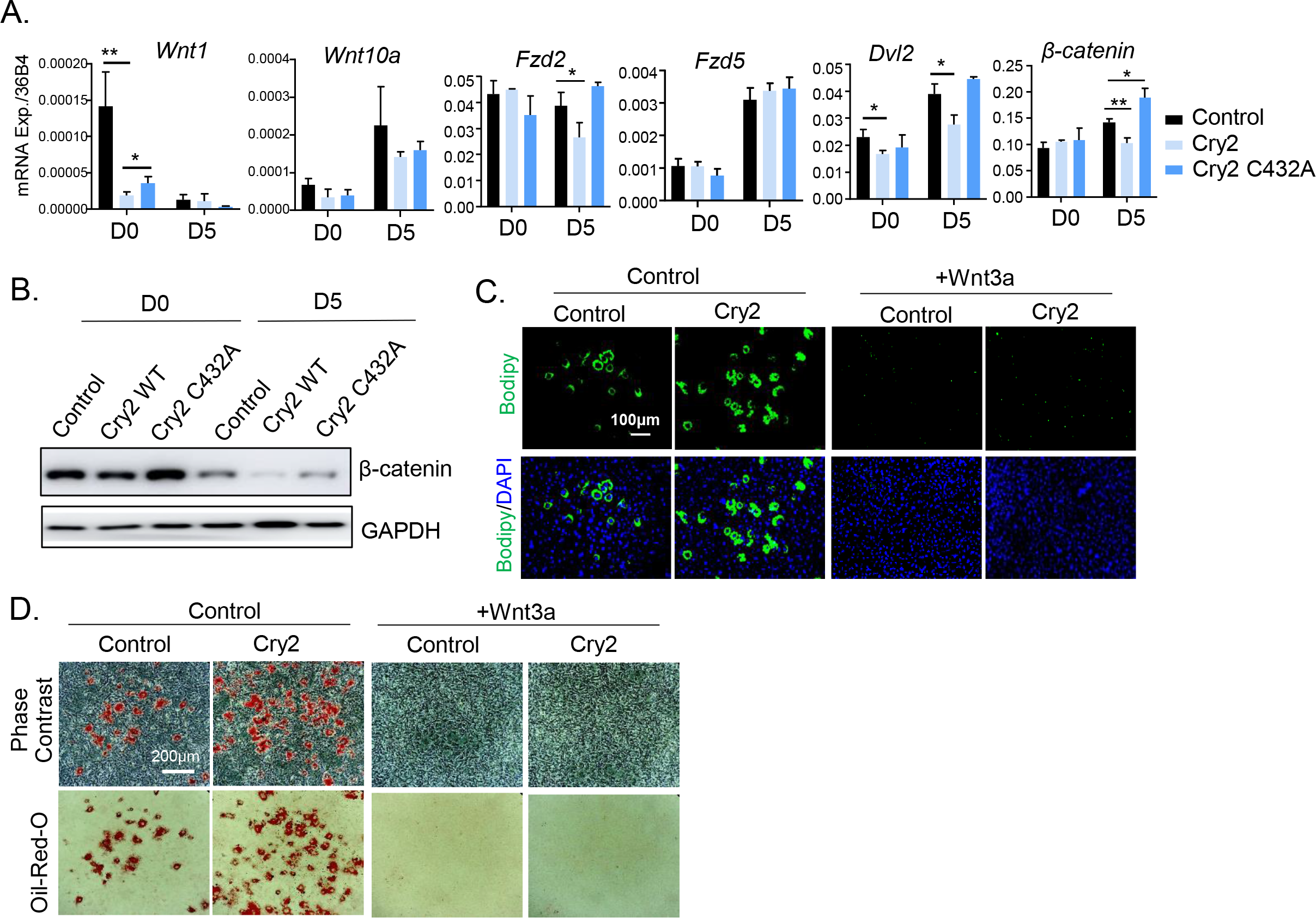
Cry2 transcriptional repression of Wnt signaling pathway mediates adipogenesis. (A) RT-qPCR analysis of Wnt signaling pathway gene expression at day 0 and 5 of adipogenic differentiation of 3T3-L1 with ectopic expression of Cry2 or C432 mutant. n=3. (B) Immunoblot analysis of β-catenin protein level at Day 0 and 5 of adipogenic differentiation in 3T3-L1 with ectopic expression of Cry2 or C432 mutant. (C, D) Representative images of Bodipy (C) and oil-red-O staining (D) of day 7-differentiated Cry2-expressing 3T3-L1 cells treated with or without 20% Wnt3a media.

#### Cry2 loss-of-function inhibits while stabilization by KL001 promotes adipogenesis via modulating Wnt activity

Based on findings from the repression-defective C432A mutant, we further tested whether genetic inhibition of Cry2 by shRNA silencing modulates adipocyte development. Two 3T3-L1 clones with stable Cry2 knockdown were generated via lentiviral transduction, with both achieved ∼70% reduction of Cry2 transcript (Fig. 5A). When these cells were subjected to adipogenic differentiation, compared to cells with vector control that displayed robust differentiation after 6 days, the shCry2 clones demonstrated markedly impaired mature adipocyte formation as indicated by oil-red-O staining (Fig. 5B). Similar effects of attenuated adipocyte maturation were observed by Bodipy staining in these cells with Cry2 inhibition (Fig. 5C). Analysis of adipogenic gene induction revealed the lack of mature adipocyte marker FABP4 expression at day 3 and day 6 of differentiation in cells with Cry2 silencing as compared to its marked induction in controls (Fig. 5D). Furthermore, using the Wnt-responsive TOPFlash luciferase reporter to assess Wnt signaling activity, we found that Cry2-deficient cells displayed significantly elevated activity than controls in response to Wnt3a stimulation, indicating that inhibition of Cry2 augmented Wnt activity (Fig. 5E).

**Figure 5.**
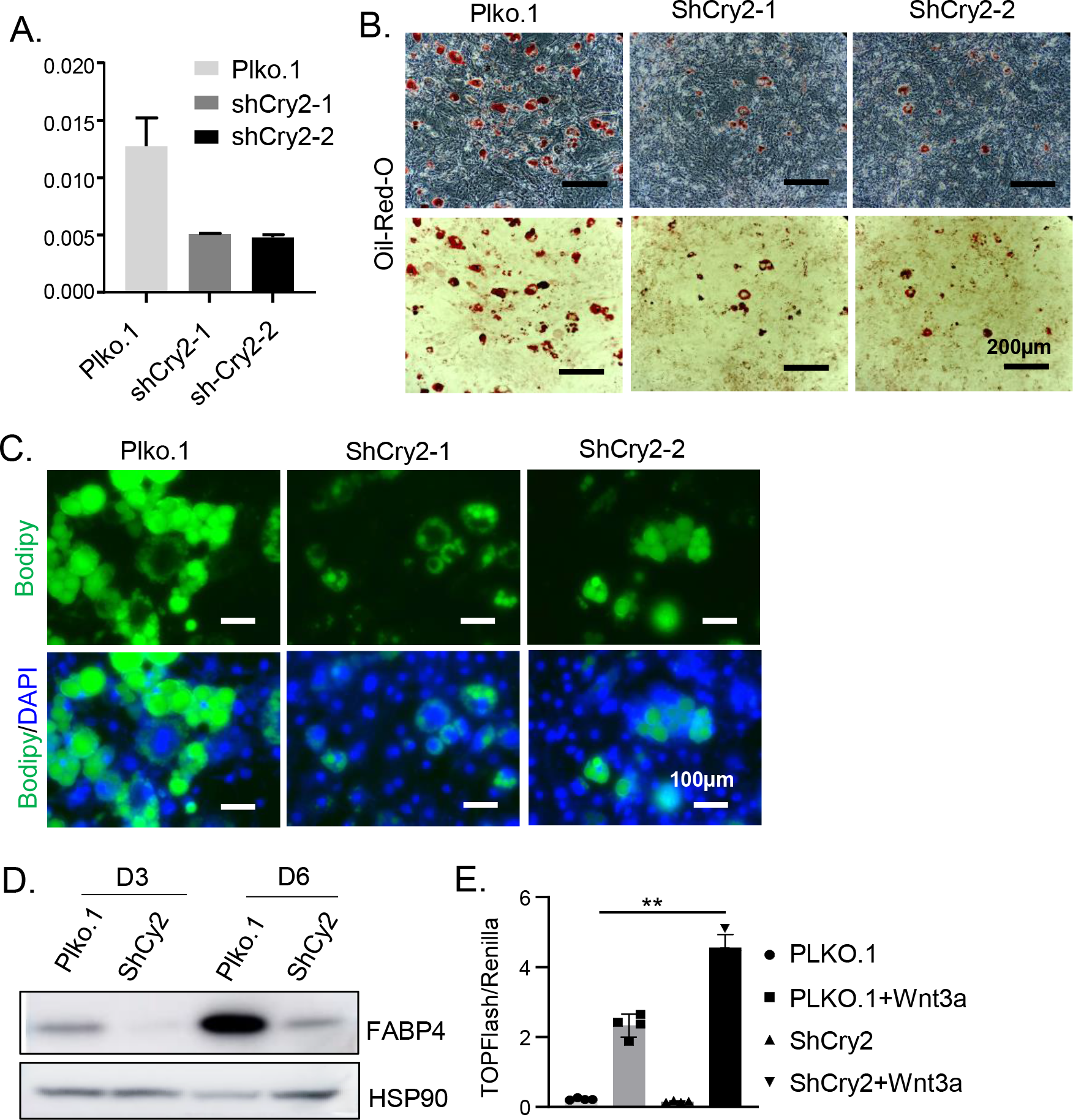
Genetic inhibition of Cry2 function impairs adipogenic differentiation. (A) RT-qPCR analysis of Cry2 transcript in 3T3-L1 clones with stable knockdown of Cry2 by shRNA, with Plko empty vector control. (B, C) Representative images of oil-red-O (B) and Bodipy staining (C) of day 7-differentiated 3T3-L1 with stable Cry2 shRNA silencing. (D) Western blot analysis of mature adipocyte marker FABP4 at day 3 and 6 of adipogenic differentiation of 3T3-L1 with Cry2 shRNA silencing. (E) Analysis of Wnt signaling activity by TOPFlash luciferase reporter in cells with stable Cry2 silencing in the presence or absence of Wnt3a stimulation. Values are represented as mean ± SD of 4 biological replicates. **: P≤0.01 by Student’s t test.

We next determined whether a pharmacological agent, KL001, which stabilizes Cry2 by blocking FBXL3-mediated protein degradation(Hirota et al. 2012), mimics that of the Cry2 gain-of-function in modulating adipogenesis. As expected, treating 3T3-L1 preadipocytes with KL001 prolonged Cry2 protein half-life, as indicated by its significantly reduced rate of degradation in the presence of cycloheximide (Fig. 6A). In line with its effect on promoting CRY2 stability, addition of KL001 augmented the Cry2 repressive activity on CLOCK/Bmal1-induced transcription activation of the Per2 promoter-driven luciferase activity (Fig. 6B). Consistent with this Cry2 repression-enhancing effect, KL001 treatment was sufficient to promote the adipogenic differentiation of 3T3-L1 preadipocytes, as indicated by Bodipy staining of mature adipocytes (Fig. 6C), oil-red-O staining (Fig. S1A) and elevated FABP4 protein expression as compared to DMSO-treated controls (Fig. 6D). Examination of Wnt pathway activity revealed KL001 effect on suppressing Wnt signaling in a dose-dependent manner under basal condition, as assessed by TOPFlash luciferase reporter (Fig. 6E). In addition, 20 µM KL001 was also able to inhibit Wnt-stimulated signaling activity. Consistent with this finding, a moderate reduction of β-catenin by KL001 treatment was observed in preadipocytes, and this effect was more evident when KL001 was administered to cells with ectopic expression of Cry2 (Fig. S1B). Notably, when tested in Cry2-deficient preadipocytes, KL001 effect on promoting adipogenesis was abolished, suggesting that this is a Cry-dependent activity of KL001 (Fig. 6F & S2A). Furthermore, treating 3T3-L1 with ectopic expression of Cry2 with KL001 revealed additive effects on adipogenesis, as indicated by markedly increased mature adipocyte formation than that of the Cry2-overexpressing cells or KL001 treatment alone of preadipocytes containing vector control using staining by oil-Red-O (Fig. 6G) and Bodipy (Fig. S2B). Analysis of adipogenic inductions via PPARγ and FABP4 further validated the additive effects of KL001 treatment and Cry2 on adipogenesis (Fig. 6H). Taken together, using distinct approaches to probe Cry2- Per2 interaction and its clock repressor function in adipogenic progenitors, we demonstrated that inhibition of Cry2 function, via a mutant defective of Per2 interaction or genetic silencing, relieves repression of CLOCK/Bmal1 activity to potentiate Wnt signaling that suppresses adipocyte development. Conversely, promoting Cry2 function, either genetically or pharmacologically, led to inhibition of CLOCK/Bmal1 transcriptional control of Wnt pathway to promote adipogenesis (Fig. 7).

**Figure 6.**
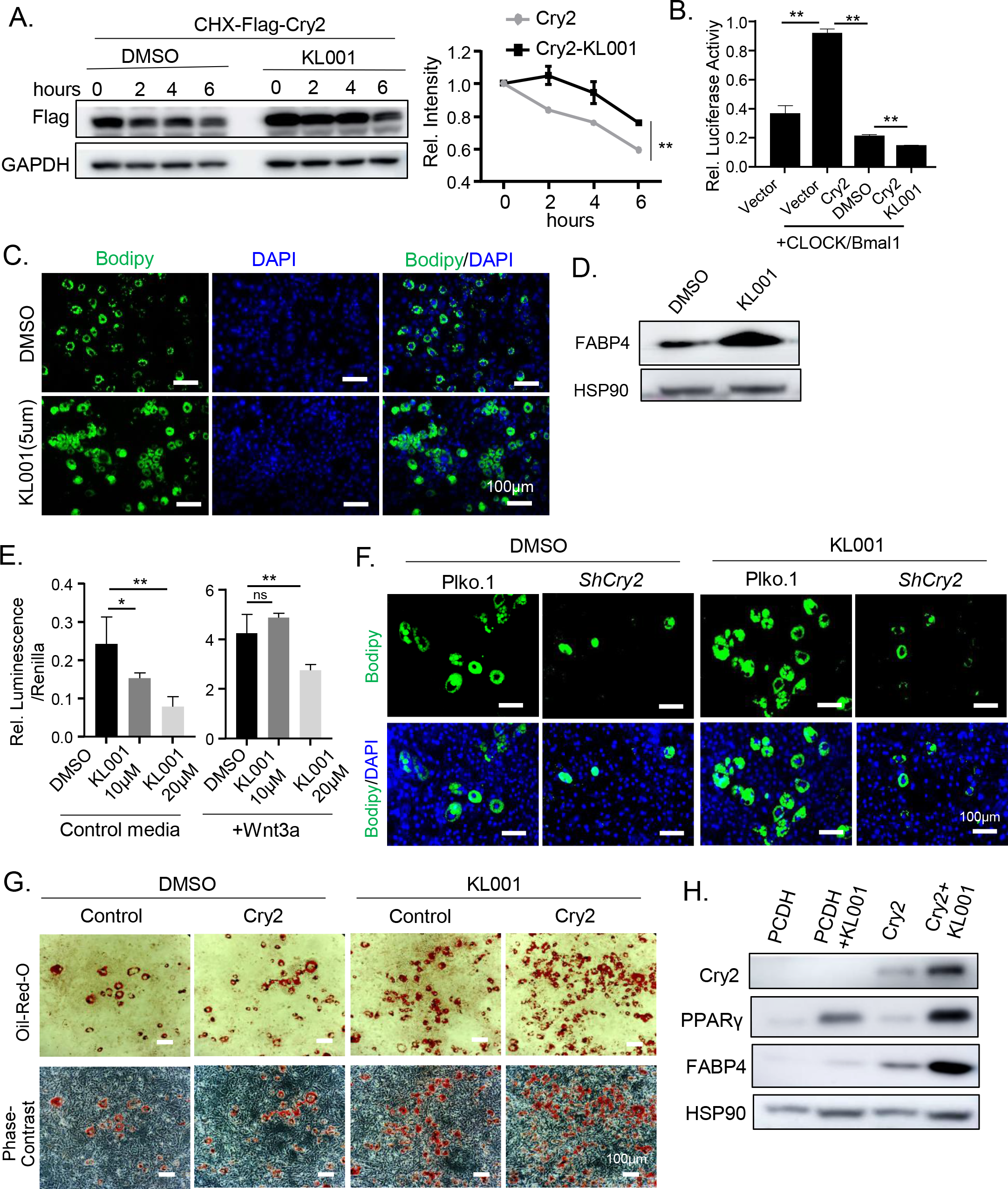
Cry2-stabilizing compound KL001 suppresses Wnt signaling to promotes adipogenesis. (A) Immunoblot analysis of KL001 effect on Cry2 protein half-life in HEK293A cells transfected with Flag-tagged Cry2 with the presence of cycloheximide. Cells were treated with DMSO or KL001(10 µM) in the presence of CHX (50 ng/mL) for 8 hours at 24h following transfection of Flag-tagged Cry2. Densitometry was performed with normalization to GAPDH for quantitative analysis of Cry2 protein abundance. **: P≤0.01 by one-way ANOVA. (B) The effect of KL001 on transcriptional repression activity of Cry2 or mutant on CLOCK/Bmal1-mediated activation of a *Per2:dLuc* reporter. HEK293A cells were transfected with Per2-luc, Bmal1 and CLOCK with Cry2 as indicated. At 24 hours following transfection, cells were treated with KL001(10 μM) or DMSO for 8 hours prior to analysis of luciferase activity. **: P≤0.01 by Student’s t test. (C) Representative images of Bodipy staining of 3T3-L1 at day7 of differentiation with or without KL001 treatment. KL001 (5 µM) was added at induction of differentiation. (D) Immunoblot analysis of mature adipocyte marker FABP4 expression at day 7 of 3T3- L1 differentiation with or without KL001 treatment. (E) Analysis of KL001effect on Wnt signaling activity using TOPFlash luciferase reporter in 3T3-L1 preadipocytes with or without 20% Wnt3a stimulation. Values are represented as mean ± SD of 4 biological replicates. (F) Representative images of Bodipy staining at day 7 of differentiation of 3T3-L1 with Cry2 silencing with or without KL001 (5 µM) treatment. (G, H) Representative images of oil-red-O staining (G) and immunoblot analysis of adipogenic differentiation genes (H) of day 7-differntiated Cry2-expressing 3T3-L1 cells with or without KL001 (5 µM) treatment.

**Figure 7.**
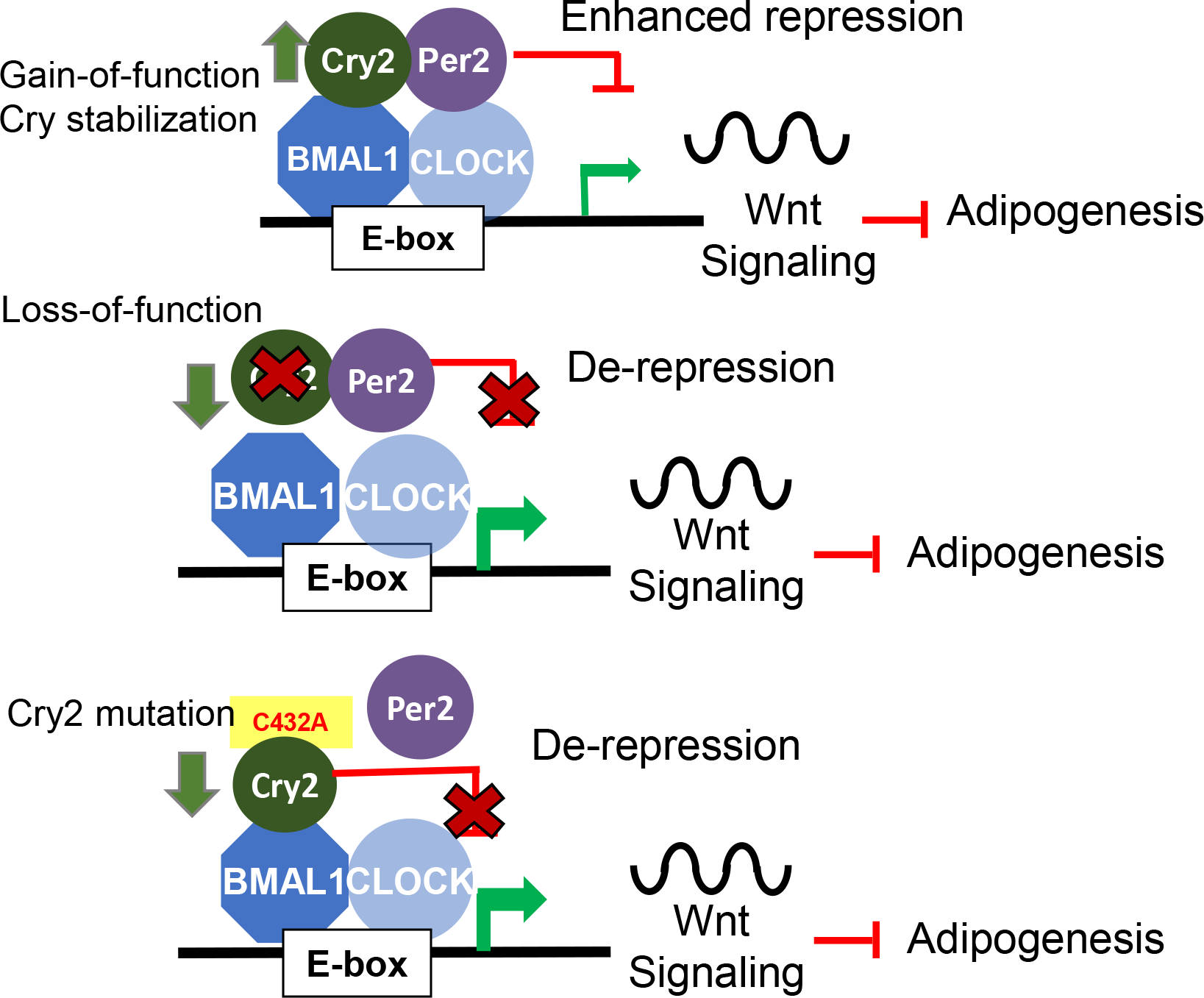
Proposed model of Cry2-Per2 transcription repression in modulating clock control of Wnt signaling that promotes adipogenesis.

## Discussion

Despite studies indicating an important function of Cry2 in metabolic regulations and the role of circadian clock in orchestrating adipogenic processes, how Cry2 and its interaction with Per2 may modulate adipocyte development remains largely unknown. Employing distinct approaches to interrogate Cry2 repressor function, *via* a point mutation that abolished CRY2-PER2 interaction together with genetic and pharmacological tools, our study revealed the activity of Cry2-mediated repression in promoting adipogenic maturation through transcriptional control of the Wnt signaling pathway.

Based on the structural analysis of Per2 Cry-binding domain and Cry2 photolyase homology region (PHR) complex, Per2 encompass a broad surface that interacts with C-terminal region of the FAD- binding domain of Cry2 (Nangle et al. 2014; Schmalen et al. 2014). This interaction competes with FBXL3 binding, thus preventing CRY from proteasome-mediated degradation. KL001 binds within the CRY FAD-binding domain, thus stabilizing CRY protein by blocking access of FBXL3. C432 lies within an interface loop structure of Cry2 close to the coil-coil domain that forms an intermolecular zinc finger with Per2 (Nangle et al. 2014). This structure is critical for PER2-CRY interaction required for the nuclear translocation and CRY repression of CLOCK/BMAL1 activation of transcription. Our results of the C432 mutant identified that this residue is a key site within the interface loop that mediates CRY2-PER2 complex formation while maintaining the ability to interact with Bmal1. As a result of abolished PER2 binding, C432A mutant displayed significant loss of repressive activity on CLOCK/BMAL1-induced transcription, indicating this is a loss-of-function mutation. Interestingly, in comparison, mutation of C430 within its proximity enhanced Cry2 repression, suggesting the specificity of these distinct residues in modulating Cry2 activity. Consistent with the finding of loss of repression on clock transcription, when tested for its effect on modulating adipogenesis, the C432A mutant showed diminished effect on promoting mature adipocyte development as compared with the wild-type CRY2. Furthermore, genetic inhibition of CRY2 led to attenuated adipogenic differentiation, providing additional support of the positive regulation of Cry2 on adipogenesis. Thus, disrupting PER2-CRY2 interaction resulted in comparable outcome as genetic CRY2 loss-of-function due to the loss of CRY2 transcription repression. On the other hand, enhancing CRY2 protein stability, and thus, its repression on CLOCK/BMAL-mediated clock activation, KL001 exerted similar effect as CRY2 ectopic expression that augmented adipogenic differentiation.

Our findings of CRY2 and KL001 modulation of adipocyte maturation adds to our understanding of the role of positive and negative arms of core clock loop in orchestrating temporal control in adipogenesis. Transcription repression by CRY2, which inhibits CLOCK/BMAL1 activity in inducing clock and clock-controlled output pathways, suppresses key components involved in the Wnt signaling cascade. A study of genome-wide Bmal1 chromatin binding sites identified its direct transcriptional control of the Wnt pathway (Rey et al. 2011), and we demonstrated previously that Bmal1 functional regulation of Wnt signaling activity promotes myogenesis while suppressing adipocyte development (Guo et al. 2012; Chatterjee et al. 2013). The transcriptional repression of CRY2 on Wnt pathway genes is likely mediated by inhibiting CLOCK/BMAL1 activity, leading to enhanced preadipocyte differentiation. In line with current findings of CRY2 effect on promoting 3T3-L1 differentiation, loss of CRY1 and CRY2 markedly attenuated lipid accumulation in brown adipose tissue due to inhibition of brown adipocyte differentiation (Miller et al. 2020). While we specifically interrogated the role of CRY2 in adipogenic regulation, CRY1 was reported to display a similar inhibitory effect on Wnt pathway that promotes adipogenesis (Sun et al. 2018). However in contrast, the PER proteins function in distinct manners in adipogenic regulations, as Per2 and Per3 were reported to interact directly with adipogenic regulators such as PPARγ (Grimaldi et al. 2010) or KLF15 (Aggarwal et al. 2017) to modulate differentiation. It remains to be further explored how loss of PER function may differ from that of CRY in adipogenic processes.

In the liver, CRY1 blocks the activity of cAMP-mediated CREB activation to inhibit hepatic gluconeogenesis during fasting to feeding transition (Zhang EE et al. 2010). A recent study indicated that impaired GSK-3β activity in stimulating Cry1 degradation contributes to elevated glucose level in diabetic animals due to uncontrolled glucose production from the liver (Kim et al. 2022). In addition, genetic ablations of Cry1 and Cry2 in mice led to elevated glucose level (Lamia et al. 2011). KL001 and related derivatives, due to CRY-stabilizing activities, demonstrated efficacy in ameliorating hyperlipidemia in diabetic models, which was attributed largely to their efficacy on inhibiting hepatic gluconeogenesis to prevent glucose production in the liver (Hirota et al. 2012; Miller et al. 2020). Based on KL001 effect on potentiating Cry2-mediated repression mechanism, we uncovered its novel activity in promoting adipogenic differentiation. As mature adipocyte is a major insulin-sensitive cell type in mediating glucose disposal, it will be intriguing to explore whether KL001, or its derivatives, may modulate adipocyte development *in vivo* and contribute to insulin sensitivity. Thus, potential tissue-specific mechanism of actions of Cry2-modulating molecules in modulating glucose metabolism remains to be dissected (Hirota et al. 2012; Miller et al. 2020).

Insulin-stimulated lipid storage in mature adipocytes provides a major mechanism for energy reserve during periods of nutrient availability. In the liver, SREBP1c activation during feeding or by insulin stimulation led to induction of CRY1 (Jang et al. 2016). Given the established role of CRY in suppressing CREB-mediated signaling in gluconeogenesis to promote the metabolic switch from a fasting to a fed state (Zhang EE et al. 2010), CRY2 activity in adipogenesis could be a coordinated response to nutrient oscillation for transitioning to feeding by promoting adipocyte lipid storage. The CRY2 activity in adipose tissue may function as a mechanism to orchestrate nutrient partitioning during the fasting to feeding metabolic adaptation, which warrants further studies *in vivo*. To date, studies of Cry2 genetic loss-of-function are confined to global ablation (van der Horst et al. 1999; Barclay et al. 2013), which precludes the dissection of its tissue-specific metabolic roles in whole-body metabolic homeostasis. Tissue-selective genetic models are needed to address Cry2 function in adipose depots and elucidate whether its dysregulation impacts obesity and glucose homeostasis. In light of the recent identification of a Cry2 missense mutation in humans that led to advanced sleep phase (Hirano et al. 2016), metabolic alterations due to CRY2 dysfunction could contribute to the development of metabolic disorders with clinical implications. Particularly, as circadian misalignment predisposes to the development of insulin resistance and obesity (Pan et al. 2011; McHill et al. 2014), pharmacological modulation of CRY2 activity in adipocyte may lead to the discovery of novel targeted clock interventions for metabolic diseases.

## Acknowledgements

We thank Drs. Seung-Hee Yoo at The University of Texas Health Science Center at Houston, Steve Kay and Meng Qu at the University of Southern California for sharing the luciferase reporter plasmids and cell lines used in this study. KM is a faculty member supported by the NCI-designated Comprehensive Cancer Center at the City of Hope National Cancer Center. This project was supported by a grant from National Institute of Health 1R01DK112794 and a Type II Diabetes Innovation Award from Arthur Riggs Diabetes and Metabolism research Institute to KM. The funders had no role in study design, data collection and analysis, decision to publish, or preparation of the manuscript.

## Authorship Statement

WL, XX and TL: data curation and investigation, formal analysis, manuscript editing; KM: formal analysis, project administration, manuscript writing and editing, and funding acquisition.

## Data Availability Statement

Data that support the findings of this study, including the full set of images of raw immunoblot data and stainings presented, will be available via Mendeley data repository through accessing a link when deposit is completed.

