## Supplemental Figures for "Transcription repression of Cry2 via Per2 interaction promotes adipogenesis"

### Supplemental Figure S1.

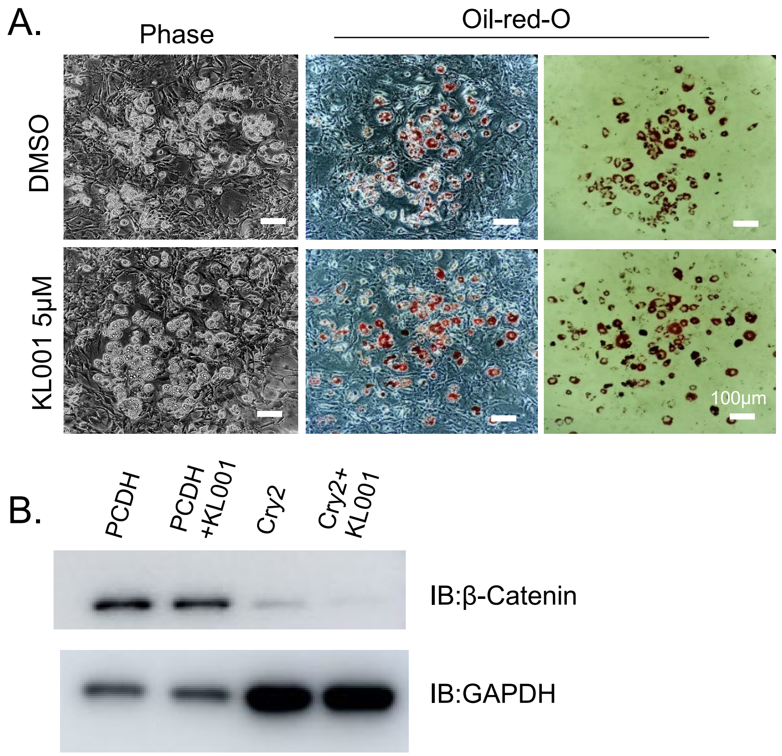


**Figure S1. The effect of Cry-stabilizing compound KL001 on promoting adipogenesis.**

(A) Representative images of phase contrast and oil-red-O staining of day 6-differentiated 3T3-L1 treated with DMSO or 5μM of KL001. (B) Western blot analysis of β-catenin protein expression in differentiated control or Cry2-overexpressing 3T3-L1 treated with or without KL001 treatment.

### Supplemental Figure S2.

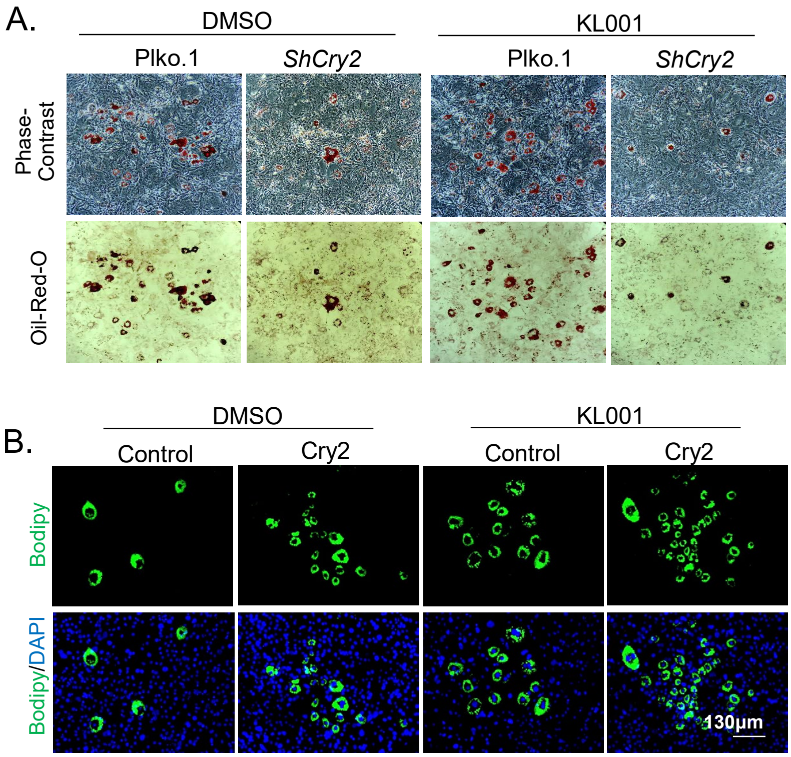


**Figure S2. The effect of KL001 on adipgenesis in 3T3-L1 cells with genetic silencing or ectopic expression of Cry2.**

(A) Representative images of oil-red-O staining of day 6-differentiated 3T3-L1 containing vector control (Plko.1) or Cry2 shRNA (shCry2), treated with DMSO or 5μM of KL001. (B) Representative images of Bodipy staining of 6-differentiated 3T3-L1 containing vector control or Cry2-overexpressing treated with DMSO or 5μM of KL001.

**Table S1. Primer sequence for site-directed mutagenesis of Cry2.**

| | |
|---|---|
| Cry2C430A-s | ATTCTTCCACGCCTACTGCCCT |
| Cry2C430A-as | GCGTGGAAGAATTGTTGGAAGAAAGCA |
| Cry2C432A-s | CCACTGCTACGCCCCTGTGGGC |
| Cry2C432A-as | GCGTAGCAGTGGAAGAATTGTTGGAAG |
| Cry2C430432A-s | TTCTTCCACGCCTACGCCCCTGTGGGC |
| Cry2C430432A-as | GCGTAGGCGTGGAAGAATTGTTGGAAG |

### Table S2. Primer sequences for Cry2 shRNA.

| | |
|---|---|
| ShmCry2-1 | GCTCAACATTGAACGAATGAA |
| ShmCry2-2 | GGTTCTCTTCACAAGGTATAA |

### Table S3. List of primary antibodies.

| Antibody | Source | Cat# |
|---|---|---|
| Cry2 | Thermo Fisher | PIPA579073 |
| Flag-Tag | Sigma Aldrich | F7425 |
| HA-Tag | Cell Signaling | 3724S |
| Myc-Tag | Cell Signaling | 2278s |
| β-Catenin | Cell Signaling | 8480S |
| P-β-Catenin | Cell Signaling | 4176S |
| Bmal1 | Cell Signaling | 14020S |
| Per2 | Proteintech | 67513-1-IG |
| CEBPα | Santa Cruz | SC-61 |
| PPARγ | Santa Cruz | SC-204 |
| FABP4 | Santa Cruz | SC-27159 |
| HSP90 | Cell Signaling | 4874S |
| GADPH | Thermo Fisher | AM4300 |

**Table S4.** Sequences of RT-qPCR primers.

| | | |
|---|---|---|
| 36B4 | Forward | CGCTTTCTGGAGGGTGTCCGC |
| | Reverse | TGCCAGGACGCGCTTGTACC |
| Cry1 | Forward | CTGGCGTGGAAGTCATCGT |
| | Reverse | CTGTCCGCCATTGAGTTCTATG |
| Cry2 | Forward | TGTCCCTTCCTGTGTGGAAGA |
| | Reverse | GCTCCCAGCTTGGCTTGA |
| Bmal1 | Forward | CGCTTTCTGGAGGGTGTCCGC |
| | Reverse | TGCCAGGACGCGCTTGTACC |
| CLOCK | Forward | TTGCTCCACGGGAATCCTT |
| | Reverse | GGAGGGAAAGTGCTCTGTTGTAG |
| β-Catenin | Reverse | CGCTTGGCTGAACCATCAC |
| | Forward | GTTCCGCGTCATCCTGATAGT |
| DVL2 | Forward | TCAGTTTGCGGGTGTGCGCAG |
| | Reverse | TTCGTCTCGCCTACACCACCG |
| TCF3 | Forward | AGGGCCTGCCAGGGACATCA |
| | Reverse | GGATGGCCTCGTCCAAGCGG |
| AXIN2 | Forward | TGACTCTCCTTCCAGATCCCA |
| | Reverse | TGCCCACACTAGGCTGACA |
| Fzd2 | Forward | CATGCCCAACCTTCTTGGC |
| | Reverse | CAGCGGGTAGAACTGATGCAC |
| Fzd5 | Forward | AATCATGCAGGGGGCCCCGAA |
| | Reverse | CGACAAGCTAGGTACCTGTGGCG |
| Wnt1 | Forward | GGTTTCTACTACGTTGCTACTGG |
| | Reverse | GGAATCCGTCAACAGGTTCGT |
| Wnt10b | Forward | GAAGGGTAGTGGTGAGCAAGA |
| | Reverse | GGTTACAGCCACCCCATTCC |
| PPARγ | Forward | ACCGCCCAGGCTTGCTGAAC |
| | Reverse | TGGAGCACCTTGGCGAACAGC |
| FABP4 | Forward | AAGGTGAAGAGCATCATAACCCT |
| | Reverse | TCACGCCTTTCATAACACATTCC |
| CEBPα | Forward | GGTACGGCGGGAACGCAACA |
| | Reverse | CGGCTCAGCTGTTCCACCCG |
| CEBPβ | Forward | CAAGAGCCGCGACAAGGCCA |
| | Reverse | CTCGCGACAGCTGCTCCACC |
